# Don’t bet against the odds! Odd prey in mixed species groups suffer fewer attacks than lone individuals

**DOI:** 10.1101/2024.09.02.610765

**Authors:** Akanksha Shah, Mike M. Webster

## Abstract

Mixed-species groups are common in nature. Such groups are characterised by the presence of one or more majority species, and smaller numbers of minority species. Minority individuals are expected to be subject to oddity effects; by looking or behaving differently to majority members they should be disproportionately targeted by predators. Given this, why might minority species remain in mixed-species groups? To address this question, we used threespine sticklebacks (*Gasterosteus aculeatus*) as predators and two ‘species’ of virtual prey presented via videos. We compared predator attacks on solitary prey, and odd and majority grouped prey individuals in groups of different sizes. We found that solitary prey were attacked significantly more than odd and majority grouped prey, while, in fact, odd and majority grouped prey did not differ from each other in terms of attacks received. We also found that prey in smaller groups suffered significantly more attacks than prey in larger groups. These findings provide no evidence for oddity effects but suggest evidence of a confusion effect. Natural mixed-species groups persist for various reasons, for example as foraging guilds, or because some members take advantage of more effective vigilance or alarm calls of others. We suggest, based on these findings, an additional non-mutually exclusive reason; under some circumstances, odd individuals might join larger heterospecific groups because any costs of being odd are greatly outweighed by the predation risk costs of remaining alone.

## INTRODUCTION

Swarms of insects, shoals of fish, flocks of birds and herds of ungulates: grouping is a dynamic and widespread natural phenomenon. This gregariousness in animals must persist and be favoured by selection if individuals are better able to enhance their fitness when interacting with other individuals than when alone (Landeau & Terborgh 1986). In fact, grouping has many advantages, including increased foraging efficiency, increased access to mates, resource conservation and improved navigation. For many animals, the main benefit of grouping is related to reducing predation risk (Krause & Ruxton 2002; Ward & Webster 2016).

Larger groups are more effective in detecting prey through collective vigilance (Cresswell 1994; Dill & Ydenberg 1987; Foster & Treherne 1981). The ‘many-eyes’ effect (Pulliam 1973) suggests that as group size increases, there are more individuals scanning for predators who can transmit an alarm upon detection. Hence, there is a reduced need for individual vigilance coupled with a reduction in the risk of failing to detect a potential threat. This enables members of large groups to invest time and energy in other activities.

Additionally, as the group size increases, the likelihood that any one individual will be attacked and captured is expected to decrease through dilution and attack abatement (Turner & Pitcher 1986). While larger groups may be more conspicuous to predators, each individual member of a group has a diluted risk of being attacked as the number of group members increases (Ward & Webster 2016). Wrona & Dixon (1991) reported this effect in their study on worm predation on the pupae of sedge fly (*Rhyacophila vao*). They found that although larger groups were attacked more often, through attack dilution effects, the likelihood of attacks directed to a focal individual was reduced in larger groups.

Large groups may also benefit from an enhanced predator confusion effect. The confusion effect describes the decrease in a predator’s ability to in single out and attacking a specific target when faced with a group of moving prey. This is attributed to the increased visual stimulation created by the movement of a group and the higher cognitive load of tracking an individual in such a group (Krakauer 1995). In a set of experiments combining neural network models and experimental work, Ioannou et al. (2008) found that the attack success rates of three-spined sticklebacks (*Gasterosteus aculeatus*) decreased as the group sizes of Daphnia magna increased. This was attributed to poor neural mapping in sticklebacks and increased error in targeting prey as swarms became larger. The authors also found that confusion was caused by the increased number of group members rather than prey density and area which have minimal effects on predator confusion.

Along with group size, group composition may also affect the anti-predatory advantages of grouping and shape the grouping decisions of animals (Krause & Godin, 1994). Variation in group composition can arise from differences in members’ phenotype, including colouration (Landeau & Terborgh 1986), body size (Theodorakis 1989), or ability to coordinate movement (Ioannou et al. 2012), and species (Allan & Pitcher 1986; Goodale et al. 2017). Such variation can counter the confusion effect and increase the attack success of predators (Paijmans et al. 2019). This disproportionate targeting of individuals who stand out from the rest of the group by predators is known as the oddity effect. In a system where predators hunt visually, confusion and oddity effects may be expected to favour uniformity in prey groups composition (Blakeslee et al. 2009; Krause & Godin 1994), and selection against mixed-species groups.

In fact, mixed-species groups are common in nature (Goodale et al. 2017), and are seen in fish (Allan 1986; Karplus et al. 2007; Paijmans et al. 2019), birds (Goodale et al. 2014; Satischandra et al. 2007) and mammals (Sinclair et al. 1985; Stensland et al. 2003). While forming of mixed-species groups may lead to larger overall group sizes, any dilution effects or other benefits are expected to be offset by reduced confusion and increased oddity effects.

Several studies have investigated the interaction between group size and varied composition on mixed-species groups. Fitzgibbon (1990) investigated predator attacks on mixed-species herds of Thomson’s and Grant’s gazelles (*Gazella thomsoni* and *G. granti*), reporting that cheetahs (Acinonyx jubatus) avoided attacking larger groups, but when attacks were performed, they were targeted towards the Thomson’s gazelle, who are smaller, slower, and less agile than Grant’s gazelles. Similarly, Wolf (1985) investigated the effect of predator presence on the grouping of striped parrotfish (*Scarus iserti*), stoplight parrotfish (*Sparisoma viride*) and ocean surgeon fish (*Acanthurus bahianus*), observing that when the large mixed-species groups were threatened by a predator, odd individuals in the group separated to form smaller single-species groups, perhaps to offset oddity effects.

Finally, in a classic experiment, Landeau and Terborgh (1986) investigated the influence of oddity effects on grouping, by varying the number of oddly coloured minnows in a school and measuring the attacks received by individuals in the group. It was found that although odd individuals were preferentially attacked, they still benefited from the confusion effect as they were victims of only about fifty percent of attacks. Lone fish, on the other hand, were successfully attacked in every trial. Furthermore, when one or two odd prey were added into larger schools, the presence of odd individuals did not elevate the attack rates or increase predation over the rates recorded for uniform schools.

These studies provide conflicting implications for odd individuals in mixed-species groups. While all the studies acknowledge that odd individuals experience the oddity effect to some degree, it is only Landeau and Terborgh (1986) who reported a negligible effect of oddity effects at larger group sizes. Hence, it is unclear whether odd individuals experience a disproportionate amount of targeting from predators when in mixed-species groups, especially, when these groups are larger. Additionally, if odd individuals do experience a disproportionate cost, it is unclear why these individuals should participate in such a grouping. Therefore, our study aimed to test Landeau & Terborgh’s (1986) idea that odd individuals may, in mixed-species groups, experience reduced predation risk compared to lone individuals and whether the predation risk for odd individuals changes with increasing group sizes.

To achieve these objectives, predatory fish were presented with simulated prey of different ‘species’ and group size combinations and the frequency of attacks performed by the fish towards these virtual prey was measured. Simulated models of grouped animals have been widely developed to understand the formation of groups. Studies generally link these models to functional explanations such as alignment of group members and polarised movement (Mirabet et al., 2007). However, a few studies have used simulations to assess the relationship between predation and prey grouping behaviour (Vabé & Néttestad, 1997; Wood & Ackland, 2007). Additionally, Ioannou et al. (2012) demonstrated that real-life predatory fish bluegill sunfish (*Lepomis macrochrius*) readily interact with virtual prey groups and select for coordinated movement as predicted by the confusion and oddity effects.

In the present study, virtual prey groups were created and presented to fish in the laboratory and the responses were recorded. It was predicated that (i) lone prey would be attacked more frequently than odd prey within groups as odd individuals would benefit from the confusion effect caused by grouping, (ii) odd prey would be attacked more frequently than majority group members due to the oddity effect and, (iii) the predation risk experienced by individuals would change with increasing group size since the strength of confusion effect should increase for larger groups.

## METHODS

### Test subjects

Three-spined sticklebacks were collected from the Kinnessburn stream, St Andrews UK (56.336071, -2.791089) using mesh minnow traps in June and July of 2022. We only collected fish that did not display signs of reproductive condition and that were free of injuries and parasites, with discarded fish released at the location of collection. A total of 150 fish were collected and tested. These were collected in three batches of 50 fish each, with each batch held in the laboratory for one week to acclimate, tested and then released, after which the tanks were cleaned and the next batch were brought in.

Captured fish were taken to our laboratory in the School of Biology at the University of St Andrews. Fish were housed in groups of 10 sticklebacks in separate tanks. Each tank measured 30 x 30 x 45 cm with a water depth of 25cm and was furnished with gravel and artificial vegetation for shelter and aerated through air stones (two air stones in each tank). Fish were kept under a 12:12 hours (7 am-7 pm) light: dark cycle at a temperature of 10°C (± 1 °C). Pieces of black corrugate plastic were put between each tank to ensure that the behaviour of individuals in one tank does not influence the behaviour of individuals in neighbouring tanks. A piece of black corrugated plastic was also placed on top of each tank to provide shade from direct light since sticklebacks show a preference for shaded areas (Thompson et al. 2016; Jones et al. 2019). The rear of the tank was also covered by corrugated black plastic, with the tablet computer used to present the video stimuli, slotted between this and the outer rear wall of the tank. The front of the tank was left uncovered.

All fish were left to acclimatize for seven days before trials began. Fish were fed at 10 am daily on defrosted bloodworms except on testing days, when they were fed after trials had concluded. Sticklebacks were tested in the tanks within which they were housed. On trial days, artificial vegetation and air stones were removed and a tablet computer was placed on the far side of the tank where virtual prey presentations were made (Figure 1). Used fish were released at a releasing point along the bank of the Kinnessburn stream (56.336489, - 2.792975). The releasing point was further upstream to the sampling point to reduce the chances of the same individuals being sampled again.

**Figure 1:**
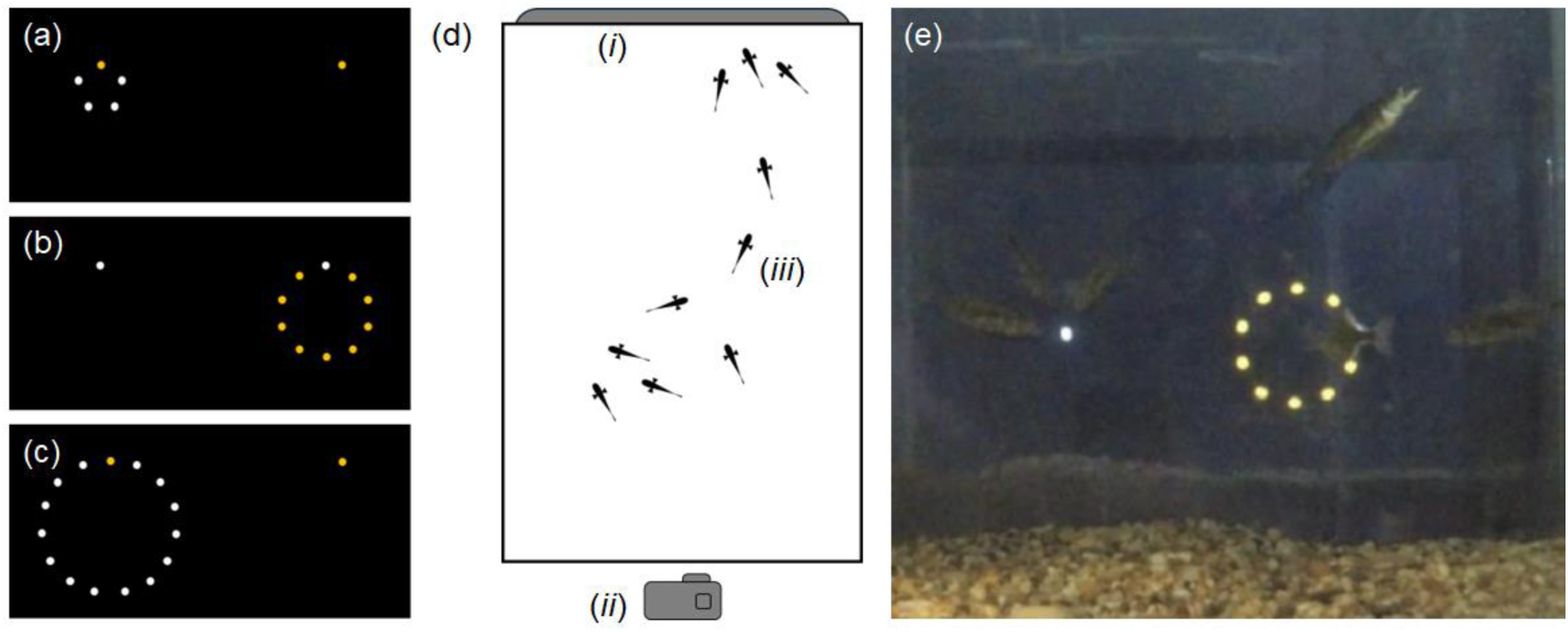
The simulations presented to predatory fish as virtual prey: (a) A solitary yellow virtual prey positioned on the right and a group of 5 consisting of 4 majority white prey and one odd yellow individual positioned on the left. (b) A solitary white prey positioned on the left with a group of 10 consisting of 9 majority yellow individuals and one odd white individual positioned on the right. (c) A solitary yellow individual on the right while a group of 15 consisting of 14 majority white individuals and one odd yellow individual positioned on the left. Note that the colour and left-right positioning of the prey were randomised between trials. (d) a plan-view diagram of the test tank. Video stimuli were presented via a tablet computer (i) placed against the outside of the tank. A GoPro camera (ii) was paced on the opposite side the tank to film the responses of the sticklebacks (iii) to the prey stimuli presented on the tablet. During trials the sides of the tank were covered with black screening and the air filter and artificial vegetation were removed. (e) A screengrab showing test subjects attacking the simulated prey.

### Experimental design: prey stimuli

To evaluate the predation risk associated with mixed grouping and solitary prey, the sticklebacks were presented with virtual prey of two phenotypes arranged in three grouping treatments (see below). The simulations were presented via an Amazon Fire tablet (model: Fire HD 10 (9th generation); screen dimensions: 25.5x 15.5 cm). The frequency of the attacks performed by the fish towards the virtual prey was measured and compared to determine the predation risk associated with the different phenotypes and group sizes.

All simulations were made using PowerPoint (Microsoft 365, version 2206). For each trial, a separate presentation containing all the different prey phenotype and grouping combinations was created. The presentation was developed such that there was a blank black slide in the beginning to give fish time to habituate to the presence of the tablet. Following this slide, the presentation was made of the treatment slides containing the different prey groups separated by blank slides with no stimuli. A blank slide was inserted after each treatment slide to provide a rest period of 48 minutes before the next treatment was presented. The order in which the treatments appeared was randomised. Each slide was timed to appear for 5 minutes and recorded to create a video file which was presented to the fish.

The virtual prey were modelled to resemble larger zooplankton. They were created to appear as dots arranged in varying groups sizes (Figure 1). The dots were created by inserting circle shapes onto slides. These were sized to be 5mm in diameter when presented on the screen. Once inserted onto the slides, the dots were arranged in a circle with each dot being 1 cm apart. Two species or phenotypes of virtual prey were created by colouring the dots white (PowerPoint colour code: #FFFFFF) or yellow (PowerPoint colour code: #FFC000). These colours were chosen as threespine sticklebacks have tetrachromatic colour vision (Bowmaker, 1998) and can perceive both colours. To control for any colour biases, half the trials were conducted where the odd prey was white and the majority yellow, with this reversed in the other half of the trials. Slides were set to have a black background to ensure that both phenotypes appear equally conspicuous from the background. The circle of dots was then grouped, and a “spin” animation was added to simulate collective movement as can be seen for example in swarms of Daphnia (O’Keefe et al. 1998). The spin animation was also timed to rotate at a naturally appropriate speed of 20mm per second (O’Keefe et al 1998).

To evaluate the predatory risk associated with being solitary vs groups of different sizes, three group treatments were created. Group treatments were composed of either 5, 10 or 15 prey items, of which one was odd. These groups and solitary prey were presented together with groups appearing on one side of the screen and solitary prey appearing on the other side (Figure 1). These were positioned so that there was a minimum 8.5 cm between the solitary prey and the group. The side on which the group and solitary prey appeared was randomised. Hence the predatory fish had a choice between attacking the solitary prey or group of 5 (4 majority, 1 odd), group 10 (9 majority, 1 odd), group of 15 (14 majority, 1 odd), where the solitary and odd prey were the same colour.

### Experimental design: treatments

Prey stimuli were presented to each group of fish via a tablet that was placed on one side of the tank. A camera (GoPro Hero 5, 1080p, 30 fps) was placed on the other side to record all the responses of the sticklebacks towards the virtual prey (Figure 1). Each group of fish was presented with 6 stimulus videos in a randomised order. These were the three group size treatments, two times each, with prey colours switched between odd and majority prey, with 48 minutes between each presentation. A total of 90 stimulations were presented to the 15 groups of fish.

The video footage for each tank was reviewed and the prey type attacked (solitary, odd or majority), and the number of attacks (hereinafter referred to as frequency of attacks) received by each prey type was measured. An attack was defined as chasing and pecking a solitary prey item or a single member of the grouped prey. The target could be reliably identified from the videos. Since individual sticklebacks could not be identified from the video footage, the frequency of attacks was measured per tank rather than per individual fish and each group (tank) of fish was defined as a separate replicate. As there were multiple majority prey in each grouping treatment, the frequency of attacks received by majority prey types was averaged to be comparable with the frequency of attacks received by the solitary and odd individuals.

### Ethics

All procedures in this study were approved by the University of St Andrews, School of Biology, Animal Welfare and Ethics Committee.

### STRANGE statement

The STRANGE framework encourages researchers to consider sampling biases in subject test pools and any implications for the wider interpretation of their findings (Webster & Rutz, 2020). Here we have used virtual prey to make inferences about the pressures acting upon real prey animals. We argue that this is a reasonable approach, first because mixed species groups composed of a majority of one species and a smaller minority of another species are well documented (Krause et al. 1996; Hoare et al. 2000; Blakeslee et al. 2009; Ward et al. 2018), and second because use of virtual prey is a well-established method of studying predatory hunting and targeting preferences (e.g. Ioannou et al. 2012). Our virtual prey were uniform in appearance and behaviour, and present a simplified stimulus compared to real prey animals.

The fish that we used as predators in this study were of a single species and only one population was examined. We only used non-reproductive adults. Furthermore, the fish traps used during collection have been reported to select bolder, more sociable and more active individuals (Álvarez-Quintero et al., 2021; Kressler et al., 2021), which may make the study sample less representative of the wider population of sticklebacks. Finally, as reported below, we saw a strong habituation effect, with attack rates on simulated prey decreasing in later stimulus video presentations. These factors should be considered when interpreting the findings of our research.

### Statistical analysis

A Generalised Linear Mixed Model (GLMM) was fitted to the data on the frequency of attacks performed by sticklebacks towards the virtual prey, using a Poisson error distribution for counts and an inverse link function. The response variable was the frequency of attacks performed by the sticklebacks. The fixed effects included two categorical co-variates and an interaction term: the prey type (solitary, odd or majority), grouping treatment (5, 10 or 15 individuals) and the interaction between prey types and grouping treatment.

The following random effects were considered: order of presentation of the different prey stimulus videos (video order), the batch in which fish were tested (recall that fish were collected and tested in three batches, batch number), the colour of odd virtual prey individual (colour of odd prey), the specific holding tank that was used to house the test subjects (tank ID), and the side of the tank that odd prey was presented on (side of tank). The best fitting model, determined by lowest AIC score and greatest deviance explained (Table 1), was included video order, replicate number, colour of odd prey as random terms, and is reported below. All statistical tests were conducted in R version 4.2.1 (R core Team 2022) and Rstudio (RStudio Team 2020).

**Table 1.** Model selection for the Generalised Linear Mixed Model (GLMM) was fitted to the data on the frequency of attacks performed by sticklebacks towards the virtual prey. All models contained prey type (solitary, odd or majority) and prey group size treatment (5, 10 or 15 individuals) as fixed effects, and the interaction between these. The following random effects were considered: order of presentation of the different prey stimulus videos (video number), the batch in which the fish were collected (batch number), the colour of odd virtual prey individual (colour of odd prey), the specific holding tank that was used to house the test subjects (tank ID), and the side of the tank that odd prey was presented on (side of tank). The best fitting model, determined by lowest AIC score and greatest deviance explained, was the first, which included video order, replicate number, colour of odd prey as random terms.

| Model | Random effects | AIC | Deviance explained (%) | Dispersion ratio |
| --- | --- | --- | --- | --- |
| 1 | Video order, batch number, colour of odd prey | 873.4 | 76.7 | 0.998 |
| 2 | Video order, batch number, side of tank | 873.8 | 76.1 | 1.010 |
| 3 | Video order, batch number | 877.1 | 76.1 | 1.006 |
| 4 | Video order, batch number, tank ID | 879.6 | 76.1 | 1.011 |

## RESULTS

Analyses (GLMM) revealed three main findings. Our first finding was that solitary prey were attacked significantly more than any other prey type, while odd prey within groups were not attacked significantly more than majority prey (Table 2, Figure 2). Second, we found that members of groups of 5 individuals were attacked significantly more than those in groups of 10 or 15 individuals. There was no difference in the frequency of attacks received by members of groups of 10 or 15 individuals. There was also an interaction between prey type and group size. Solitary prey were attacked significantly more when presented next to a group of 5 than presented next to a group of 15. This interaction was not observed for groups of 5 and group 10 and groups of 10 compared to groups of 15. Majority prey in groups of 5 were attacked significantly more than the majority prey in groups of 10 and groups of 15, while majority prey in groups of 10 and groups of 15 did not experience a significant difference in attack frequency. The attack frequency experienced by odd prey did not change significantly between the different group treatments. Odd prey did not experience a significant change in attack frequency with increasing group size (Table 2, Figure 3).

**Figure 2.**
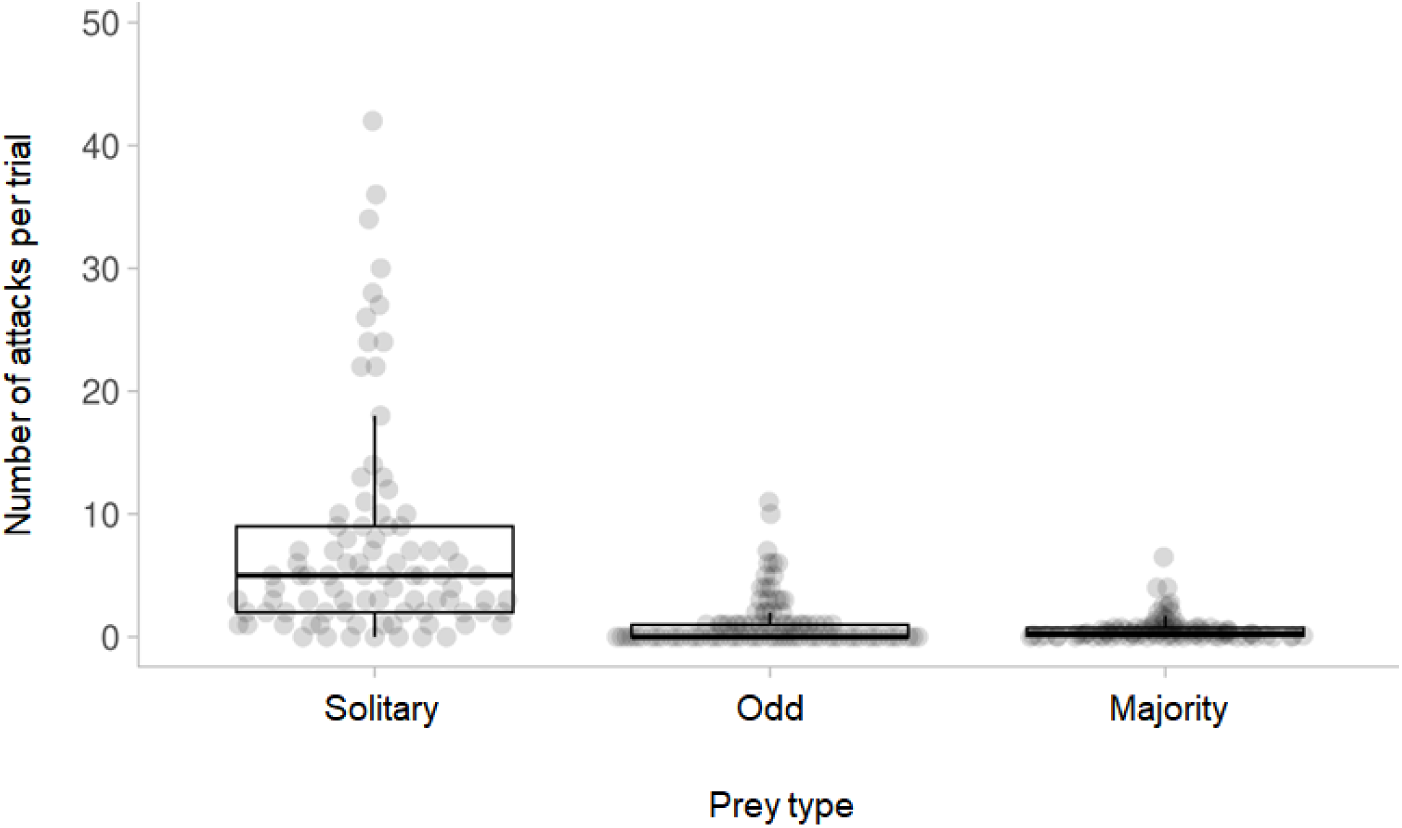
Solitary prey received significantly more attacks that did odd or majority grouped prey. Odd and majority prey did not differ in terms of number of attacks received. The bar depicts the median value, the box displays the interquartile range, the whiskers present 5 and 95 percentiles, and the dots display raw data points.

**Figure 3.**
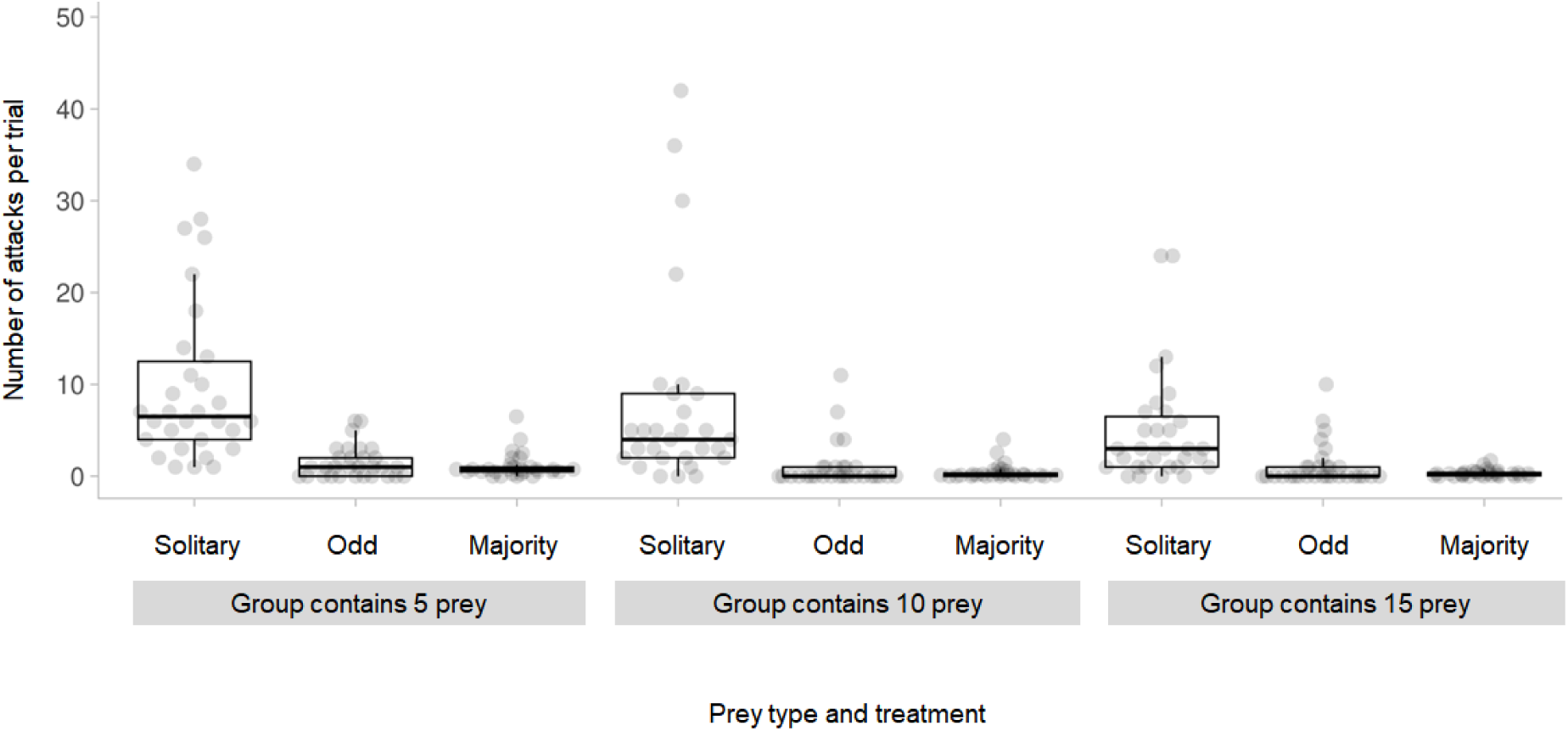
Both solitary prey that were presented alongside groups of 5 prey, and the members of groups of 5 individuals were attacked significantly more than solitary prey presented opposite, and grouped prey presented within groups of 10 or 15. There were no differences between group sizes of 10 or 15. The bar depicts the median value, the box displays the interquartile range, and the whiskers present 5 and 95 percentiles.

**Table 2.** Output of a GLMM investigating the effects of prey type (solitary prey, odd member of groups or majority member of groups) and prey group size (5, 10 or 15) upon number of attacks received. Significant effects and interactions are shown in bold. Variance for random terms is also shown.

| Term | Estimate | Standard error | Z | Pr (>\|z\|) |
| --- | --- | --- | --- | --- |
| Intercept | -0.207 | 0.385 | -0.490 | 0.591 |
| **Solitary prey vs. odd grouped prey** | **1.883** | **0.159** | **11.831** | **<0.001** |
| **Solitary prey vs. majority grouped prey** | **2.026** | **0.169** | **11.957** | **<0.001** |
| Odd grouped prey vs. majority grouped prey | 0.143 | 0.217 | 0.657 | 0.783 |
| **Group of 5 vs. Group of 10** | **-1.088** | **0.322** | **-3.376** | **<0.001** |
| **Group of 5 vs. Group of 15** | **-1.365** | **0.370** | **-3.686** | **<0.001** |
| Group of 10 vs. Group of 15 | -0.277 | 0.434 | -0.639 | 0.796 |
| Solitary with group of 5 vs solitary with group of 10 | 0.175 | 0.098 | 1.777 | 0.655 |
| **Solitary with group of 5 vs solitary with group of 15** | **0.353** | **0.102** | **3.440** | **0.014** |
| Solitary with group of 10 vs solitary with group of 15 | 0.178 | 0.107 | 1.657 | 0.736 |
| Odd within group of 5 vs odd within group of 10 | 0.270 | 0.231 | 1.169 | 0.953 |
| Odd within group of 5 vs odd within group of 15 | -0.122 | 0.213 | -0.574 | 0.999 |
| Odd within group of 10 vs odd within group of 15 | -0.392 | 0.230 | -1.706 | 0.704 |
| **Majority within group of 5 vs majority within group of 10** | **1.088** | **0.322** | **3.376** | **0.017** |
| **Majority within group of 5 vs majority within group of 15** | **1.365** | **0.370** | **3.686** | **0.006** |
| Majority within group of 10 vs majority within group of 15 | 0.277 | 0.434 | 0.639 | 0.999 |

| Random terms | Variance | Standard error |
| --- | --- | --- |
| Batch number | 0.12 | 0.35 |
| Colour of odd prey | 0.02 | 0.16 |
| Video order | 0.61 | 0.78 |

Third, the model contained batch number, video order and the colour of odd prey as random effects. Batch number and colour of the odd prey did not explain a large amount of variance, but video order accounted for a large proportion of variance (Table 2, Figure 4). Fish performed more attacks during the first video regardless of the treatment presented, and that the attack frequency decreased with successive videos, suggesting habituation to the virtual prey stimulus. Despite this effect solitary prey consistently received more attacks that grouped odd or majority prey (Figure 4).

**Figure 4.**
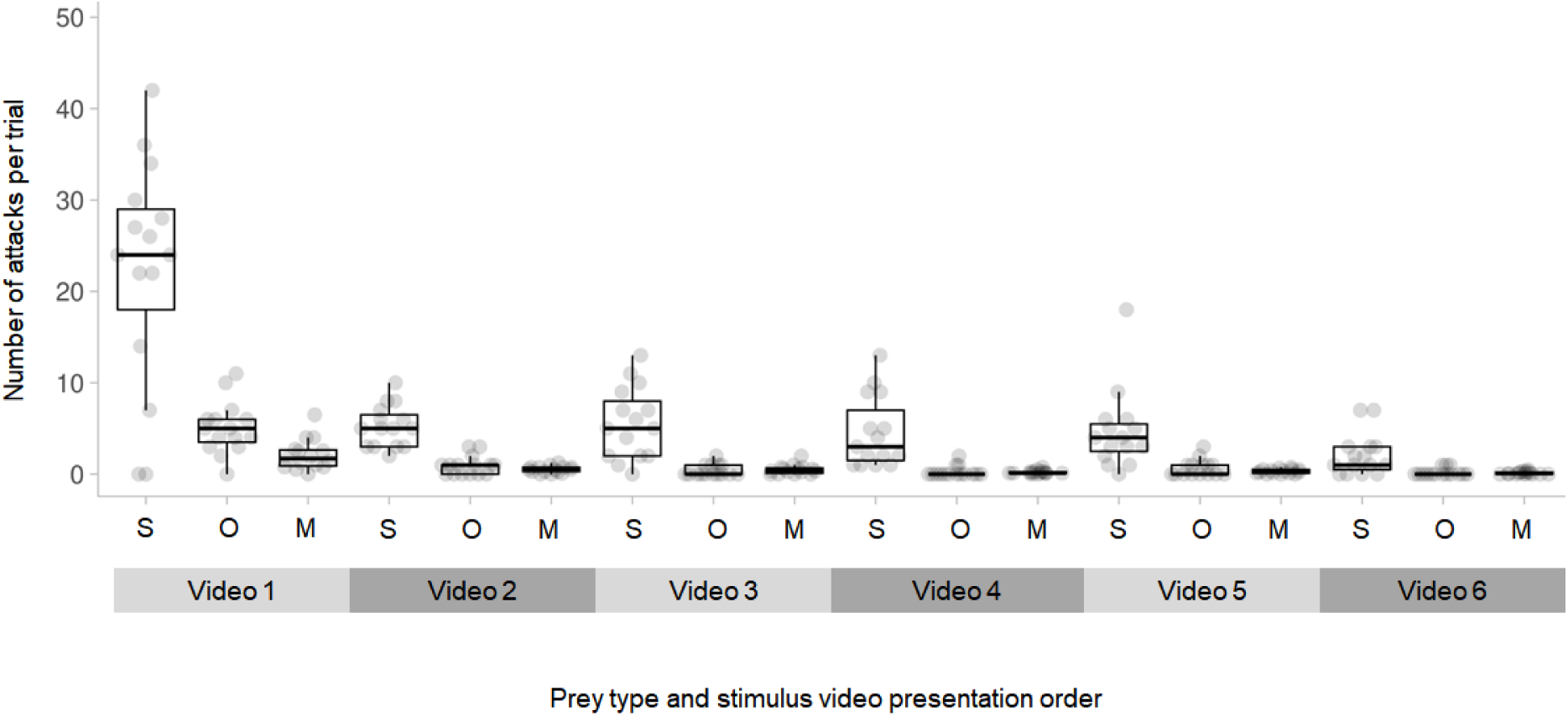
Attack frequency decreased with successive videos, regardless of stimulus presented, suggesting a habituation effect. S refers to solitary prey, O to odd members of groups and M to majority members of groups. The bar depicts the median value, the box displays the interquartile range, the whiskers present 5 and 95 percentiles, and the dots display raw data points.

## DISCUSSION

Our study provides an explanation for the persistence of mixed-species groups in nature, in the face of predation pressure. We found that solitary prey received more attacks than odd or majority prey and that prey in smaller groups suffer more attacks than prey in larger groups, supporting our first and third predictions. Our second prediction, that odd prey within groups would be attacked more frequently than majority group members, was not upheld. Instead, we saw no difference in the numbers of attacks received by odd and majority group members. Our findings are consistent with a confusion effect, that benefited odd and majority prey alike, where grouping is better than remaining alone (Landeau and Terborgh 1986) and where predator attack rates declined with increasing group size (Krakauer 1995; Ioannou et al. 2008).

Other studies have found evidence of oddity effects. Tosh et al. (2007) used a neural network model to explore confusion effects and mixed-species prey groups. The neural network was presented with groups of varying compositions of prey types and the sensory mapping of these was modelled. They found that grouping in general led to poor reconstruction in the sensory mapping of predators and that was exacerbated when prey group members were similar in appearance. Moreover, the authors observed that majority prey-types were targeted less by predators when grouped with odd individuals than when in uniform groups. In this system, odd prey provided a benefit to majority prey in that they were disproportionately less likely to be targeted when odd prey were present. Rodgers et al. (2013) provided empirical support for this effect, using a system of sticklebacks predating differently-coloured *Daphnia*, further finding that this effect was stronger when the majority species was cryptically coloured (against the background). Our study focussed upon the benefits of the majority species, while these studies suggest that mixed-species groups may also persist through benefits that disproportionately favour the majority species.

Our study only considered oddity and confusion effects, but members of mixed-species groups also benefit from other anti-predator effects, and in domains other than predation, such as foraging. For example, Dolby and Grubb (1998) investigated the benefits gained by satellite (minority) members of a mixed species winter-forming bird flocks in North America. These flocks consisted of two nuclear (majority) species, tufted titmice (*Baeolophus bicolor*) and either Carolina or black-capped chickadees (*Poecile carolinensis* or *P. articapillus*) and multiple satellite species, including downy woodpeckers (*Picoides pubsecens*) and white-breasted nuthatches (*Sitta carolinensis*). Dolby and Grubb (1998) observed than in the absence of nuclear species, the satellite species perform more vigilance-relate behaviours and appeared to perceive a higher risk of predation threat compared to when the nuclear species were present. This suggests that the satellite species may benefit from shared vigilance when forming mixed-species flocks, while the nuclear species may benefit from being in a larger group. Through reduced time invested in vigilance, all species may also gain foraging benefits from grouping together.

The combination of increased foraging and anti-predatory benefits also explains the persistence of mixed-species flocks of nuclear orange-billed babblers (*Turdoides rufescens*), and satellite ashy-headed laughingthrush (*Garrulax cinereifrons*) and greater racket-tailed drongos (*Dicrurus paradiseus*) in the lowland forests of Sri Lanka. Satischandra et al. (2007) found that drongos experienced a higher foraging success in mixed-species flocks. In these mixed-species flocks, the three species feed at different levels of the canopy, giving rise to a foraging guild of specialists, reducing the levels of competition that would occur in a single species group. Goodale and Kotagama (2005) reported that the participation of drongos benefits the other species, since drongos produce alarm calls that the other species perceive and respond to, acting a sentinel, and effectively reducing predation risk of the group as a whole.

One limitation of our study then is that it only considers predation, and not other domains, such as foraging. Here, benefits such as reduced competition for resources might offset any costs associated with oddity (where these costs exist), while competition is likely to be a drawback of forming a larger group, potentially offsetting benefits arising from confusion effects. That said, grouping is a dynamic process, and animals can form larger or smaller groups reactively, forming smaller groups when they detect food (Hoare et al. 2004) or are hungry (Riddell & Webster 2015), or larger groups when they perceive predation risk (Hoare et al. 2004).

There are other limitations associated with our study. No oddity effect was seen in our study, which may be because grouped and solitary prey were presented on the screen together. This creates a choice for the predators and, considering that tracking individuals in a group produces a high cognitive cost (Krakauer, 1995), predators may be discouraged from attacking groups and choose solitary prey more frequently instead. As a result, the test subjects might not have interacted with the grouped prey types as much as expected. This may also explain why the attack frequency experienced by odd prey did not change with group size. On the one hand this is not a problem in so far as it is ecologically valid; predators in nature that feed on swarming prey are likely to be faced with multiple differently-sized groups of prey, and lone individuals simultaneously. Targeting of lone and grouped prey by predators therefore likely occurs in this multi-group context. On the other hand, it would be valuable to be able to separate predator choice for different group sizes from confusion effects. Useful follow-up work could present solitary and grouped stimuli to test subjects separately in order to further explore this.

We also found a substantial effect of stimulus video presentation order on attack rates, with a decrease in attacks as successive videos were presented. This effect may be due to habituation. Since no food reward was obtained for attacking the virtual prey, attacks were not reinforced, and the test subjects may have lost interest in interacting with the video stimuli. Future work could overcome this by either using live prey or a food reward, by spacing the video presentation times over a longer testing period with greater intervals, or by abandoning the repeated measures approaching and using a separate group of test subjects for each stimulus video. Each approach has a downside. Live prey cannot be controlled or presented consistently, as simulated prey can, while the use of food rewards risks satiation, which could result in a similar decline in attack rates over successive trials. Using a greater inter-trial period, or using new subjects for each stimulus presentation increases the amount of time that the subjects are held in captivity and greatly increases the number of animals required, with implications for welfare. Ultimately, experimental design is a balancing act between attempting to obtain the data necessary to test a prediction, time and space constraints and the need to consider the wellbeing of the animals we study. We note here that even though attack rates declined over successive video presentations, the predators’ preference for the solitary prey remained readily apparent. We submit that our design presents a pragmatic balance between ability to detect the experimental effect of interest and a thorough discussion of the imitations of our approach (Webster & Rutz 2020).

## CONCLUSIONS

We demonstrate that odd individuals strongly benefit from grouping as they experience a significant reduction in predation risk when in groups than when alone. Odd individuals were not disproportionately targeted, instead members of smaller groups suffered higher rates of attacks, an observation consistent with a confusion effect. Thus, members of mixed-species groups, both majority and odd individuals, might benefit being members of a larger group. This study provides an explanation for the benefits gained by minority odd individuals that join mixed-species groups.

## Acknowledgements

This work was completed as part of the BL5599 Research Project module on the MSc Animal Behaviour programme by AS. We received no additional funding for this work. We thank Annie Rowe for assistance in the field.

## Author Contributions

Both authors contributed to study conceptualisation and design. AS built the equipment, ran the experiments, collected and analysed the data. AS wrote the research report and both authors edited it for publication. MMW provided general project supervision.

## Data availability

Data will be uploaded to the FigShare repository and linked to before this work is published.

